# Environmental context shapes microbial co-occurrence

**DOI:** 10.1101/2025.08.05.668733

**Authors:** Sean R. Darcy, Christina Kaiser, Ksenia Guseva

## Abstract

The analysis of microbial co-occurrence patterns promises to uncover the elusive interactions underlying the many important functions microbiomes perform. We test such co-occurrence analysis on a dataset generated by a spatially explicit meta-community generalized Lotka-Volterra model, to investigate how environmental confounders affect co-occurrence patterns. We show that strong species interactions, particularly positive ones, remain detectable despite environmental complexity. However, in this context the number of spurious co-occurrences, with no correspondence to specific ecological drivers are inflated. We also explored two types of homogenization: intrinsic (via dispersal among communities) and extrinsic (via sample aggregation). In our settings both types inflate positive co-occurrences, improving the reconstruction of the environmental signal at the cost of the precision for reconstructing interactions. Negative ones, however, exhibit contrasting responses: intrinsic homogenization obscures them, while extrinsic homogenization increases their number, but at the cost of reduced precision in linking co-occurrences to ecological drivers. Our findings underscore the complex interplay between environmental factors and sampling strategies in shaping microbial co-occurrence patterns and their ecological interpretation.

## 1 Introduction

Microbial communities perform a vast number of functions critical to human and planetary health (Rappuoli et al., 2023). Many such processes emerge from complex microbial interactions, which shape the microbial communities we find in a given time and place. The study of such interactions proves challenging as natural systems can be immensely diverse both in microbial taxa and the micro-environments they inhabit (Curtis et al., 2005; Franklin et al., 2007; Oppen et al., 2023; Vos et al., 2013). Direct observations are difficult due to the small scales at which these interactions play out (Cordero et al., 2016) and cultivation experiments are both costly and fail to account for the in situ complexity of microbial environments (Kapinusova et al., 2023). To overcome this, researchers hope to infer interactions indirectly. The idea is that interacting species either benefit or constrain each other consistently across observations. Consequently, when a pair co-occurs (or avoids each other), which is most often determined via strong correlations in their abundances (Weiss et al., 2016; Matchado et al., 2021), the pair is expected to interact positively (or negatively). This forms the foundation of microbial co-occurrence network analysis (Faust and Raes, 2012), which has seen exponential adoption in the field of microbial ecology in the past decade (Guseva et al., 2022). However, this approach is heavily criticised because interpretations of the results are often speculative, given the ambiguous origins of species co-abundance (Carr et al., 2019; Blanchet et al., 2020). This can not only result from interactions but also from species similar fundamental niche requirements (Deutschmann et al., 2021; Faust, 2021) and is likely further modified by other ecological dynamics such as dispersal and stochastic demographic events (e.g. random extinction). Furthermore, two species’ co-occurrence may arise indirectly due to influence of a third (Kurtz et al., 2015). Beyond these issues, typical studies measure relatively large volumes of material, where depending on the spatio-environmental characteristics of a system deterministic signals in co-abundance from the micro-scale might become obscured (Alteio et al., 2021; Simon et al., 2024). Despite the ample need to identify the underlying interactions that shape microbial communities we still do not understand if and to what degree the data we use is appropriate to infer microbial interaction networks. It is necessary to understand how other ecological dynamics imprint on this signal and how it is modified by the environmental characteristics of a system and the volumes at which samples are taken.

One way to improve our understanding of the utility of current analytical methods is by applying them to artificially generated data that posses key properties of natural systems. Studies have evaluated the ability of correlations to infer, or ‘reconstruct’ interactions using artificially generated data (Berry et al., 2014; Weiss et al., 2016; Pinto et al., 2022). These studies predominantly employed generalized Lotka-Volterra equations (gLVs) to create data for co-occurrence analysis. Despite certain limitations (Momeni et al., 2017; Dedrick et al., 2023), this model has shown to approximate the dynamics of microbial communities effectively (Hu et al., 2022; Venturelli et al., 2018; Angulo et al., 2019; Stein et al., 2013). But despite their utility, studies testing the inference potential of co-occurrence with gLVs have yet to account for realistic environmental dynamics. Likely the most important determinant of species abundances stems from how well their fundamental niche matches a present set of environmental factors (Hutchinson, 1957; Colwell et al., 2009; Konopka, 2009). Species with similar environmental preferences will show consistent distributions across varying environments. So far, systematic analyses have only simulated the impact of environment on co-occurrence via additions of a random values for habitat carrying capacities (Berry et al., 2014; Pinto et al., 2022). Another important yet underexplored aspect is how the spatial distribution of microbial communities influences aggregated samples, i.e. samples taken at scales that cover multiple micro-habitats. While DNA extraction and sequencing biases are reduced with larger sample volumes (Kageyama et al., 2022), theoretical studies have shown that the detection of interactions can become obscured when sampling larger scales (Armitage et al., 2019; Araújo et al., 2014). While these studies accounted for space, they neither account for dispersal nor for coherently structured habitats, which is known to significantly influence the coexistence of interacting species (Amarasekare, 2003). Recent theoretical advances with spatially explicit gLVs have demonstrated that a system’s global properties such as diversity and stability are strongly influenced by dispersal (Mattei et al., 2024; Denk et al., 2022; Garcia Lorenzana et al., 2024; Gravel et al., 2016). Despite these advances, we lack understanding on how co-occurrence is shaped when both fundamental niches and interactions affect species distributions across spatially distributed environments. In this work we address this gap by developing a gLV model, which allows us to simulate spatially explicit meta-communities with fundamental niche dynamics. Our main objectives are to uncover how shared environmental preference and interactions imprint on co-occurrence, how this is influenced by meta-community dynamics, the spatial distribution of environmental factors and the scales of sampling.

In this study we combine the most important drivers of community assembly, including species interaction, fundamental niche and dispersal dynamics. We simulate communities with gLVs in spatial habitat networks given variable environmental heterogeneity. Positive and negative co-occurrences are then assessed for their ability to reconstruct input species interactions and similar environmental preference. Simulations increase in system complexity, beginning with simple scenarios. We then include dispersal (constrained by habitat connectivity) across various scenarios of spatio-environmental heterogeneity, and analyse the effects of sample volume. Finally, we speculate on how our findings might impact the interpretation of microbial co-occurrence networks. For this, we examine network structures arising from community assembly driven by environmental dynamics or interactions alone, and how structures reorganize when both drivers act together. Overall, we offer qualitative insights for the inference of empirical co-occurrence data by covering a range of scenarios capturing the diverse properties of systems that microbial ecologists might encounter.

## 2 Material and Methods

### 2.1 Generation of cross-sectional community data

To analyse how drivers of community assembly can be reconstructed from species co-occurrence we developed a model to simulate spatially explicit community data given a range of scenarios, covering different assembly dynamics, spatial and environmental configurations as well as sampling approaches. Fig. 1 illustrates the most central elements in our model. The model was implemented in python3 and the analysis was conducted in R. All code is available in a github repository at: https://github.com/rmSeanDarcy/simulating_association. This paper is aimed at a broad audience of ecologists, so here we provide a brief overview of the model and want point readers in search of more detailed descriptions to the Supporting information (Extended Material and methods). This includes a glossary for the variables used (Tab. S1)

**Figure 1.**
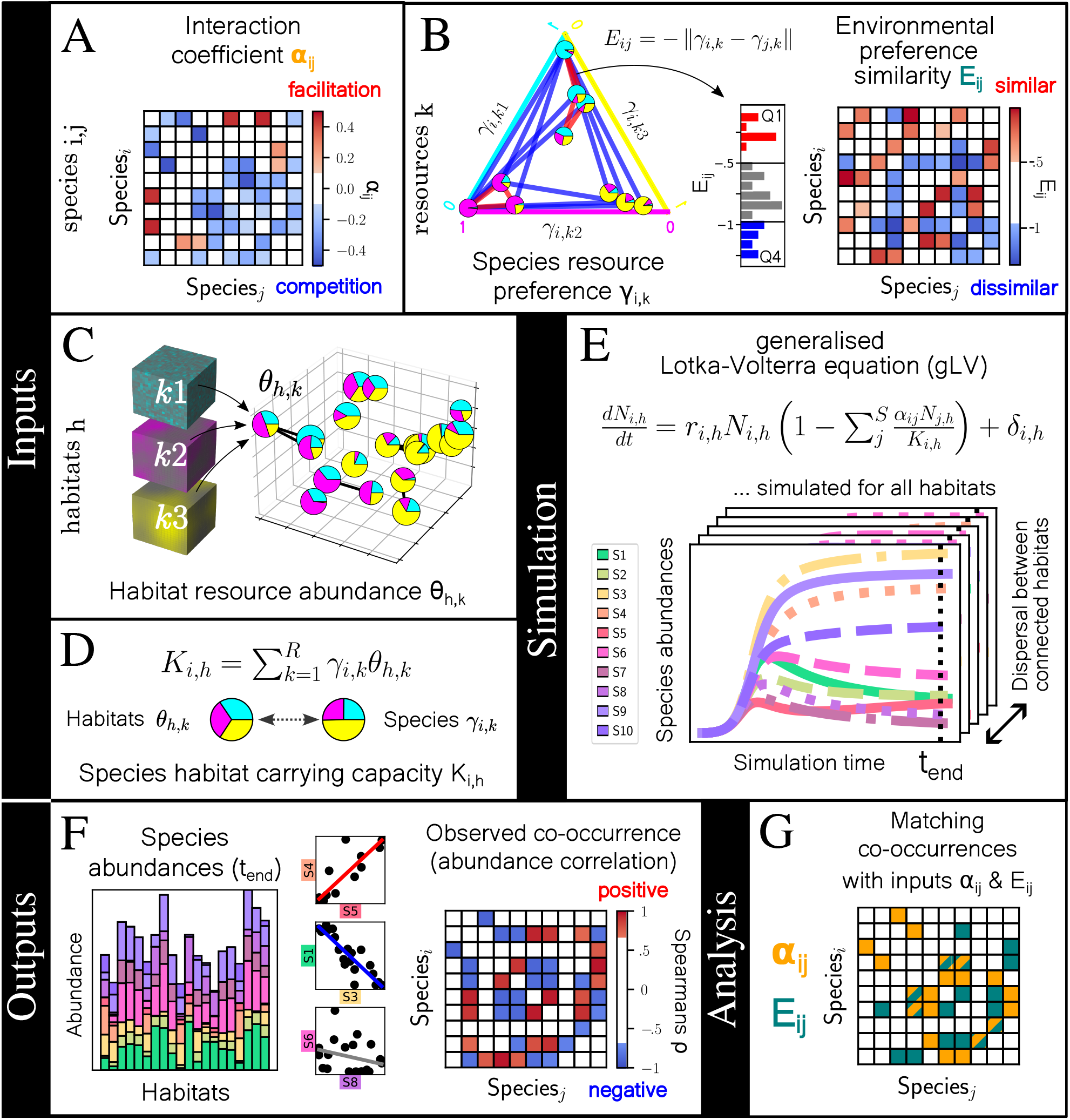
Illustrations of the most important aspects of model inputs, simulation, outputs and analysis. A) Species are generated with pairwise interaction coefficients *α*_*ij*_. 50% of all pairs either inhibit (negative = 40%) or benefit (positive = 10%) each others growth with varying strength. B) Randomly assigned preferences for three different resources *γ*_*i,k*_ produce a continuum of resource specialists and generalists. Distance between species pairs *γ*_*i,k*_ gives their environmental preference similarity *E*_*ij*_. They can be similar (first quartile, Q1), intermediate or dissimilar (Q4). C) Habitats are nodes in a spatial network. Their resource abundance *θ*_*i,k*_ are determined by projecting their coordinates into three resource landscapes of variable heterogeneity. This controls the spatial autocorrelation in habitats resource composition. Connected habitats allow dispersal. D) Species total abundance is constrained by how well their resource preferences match present resource abundances. E) Population dynamics are simulated via gLVs. Species abundances are sampled after species abundances equilibrate. F) Species co-occurrences are determined via strong positive and negative abundance correlations. G) For our main analysis, we quantify how many co-occurrences match inputs of interactions *α*_*ij*_ and similar or dissimilar environmental preference *E*_*ij*_.

#### 2.1.1 A network of habitats in spatially explicit environments

Using the networkX library (Hagberg et al., 2008), we construct random geometric graphs in which nodes represent habitats (*H*; index *h*) (Fig. 1 C). Two habitats are connected when their distance is lower or equal to the connectivity radius *d*_*e*_, allowing populations to disperse. Habitats are characterized by their position in space and values for three environmental factors (*R*; index *k*). As they quantitatively benefit species abundances we refer to these values as resource abundances *θ*_*h,k*_. We control a specific habitats *θ*_*h,k*_, by projecting its coordinates into three separate resource landscapes. These are modelled as a scale-free Gaussian Random Fields, using the FyeldGenerator library (Cadiou, 2022). The parameter *λ* controls the prevalence of small scale structures. We can thereby control the spatial autocorrelation in habitats resource composition, ranging from complete randomness (*λ* = 0), over more a fractal-like intermediate (*λ* = 5) to large patches (*λ* = 10). Every simulation therefore includes *λ* values for every *R* (*λ*_*k*1_,*λ*_*k*2_,*λ*_*k*3_). Resource distributions are further explained and depicted in the Supporting information (Extended Material and methods; Fig. S1). Resource abundances in habitats are constant, meaning they neither reduce, grow nor diffuse.

#### 2.1.2 Simulation of population dynamics

##### Generalized Lotka-Volterra and interactions

For every run we initialise species (*S* = 10) with low abundance values (*N*_*i,h*_ at *t*_0_ = 0.01) in all habitats. The biomass *N*_*i,h*_ of each species *i*, in a habitat *h* then follows our adapted generalized Lotka-Volterra differential equation (gLV):

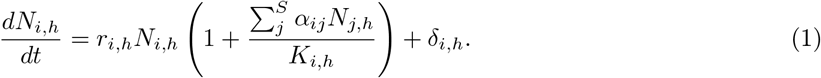

Species pairs interaction coefficient *α*_*ij*_ controls the direction (sign) and strength of an interaction, i.e. to what extent species abundances positively or negatively influence each other (Fig. 1A). Of all possible species pairs (45) about half symmetrically interact (22). Of these symmetric interactions 18 are negative and 4 are positive. *α*_*ij*_ is selected from a uniform distribution with the intervals [−0.1, −0.5] and [0.1, 0.5] respectively (see Supporting Information: Interaction coefficients). Finally, intra-specific interaction coefficients are set to *α*_*ii*_ = −1. The settings of *K*_*i,h*_, *r*_*i,h*_ and *δ*_*i,h*_ are described below.

##### Species environmental preference

For the environmental assembly dynamic we developed a species abundance is influenced by how well a habitats resource composition *θ*_*h,k*_ matches its resource preferences *γ*_*i,k*_ (Fig. 1 D). We define a species habitat carrying capacity (in Eq.1), of species *i* in habitat *h* as:

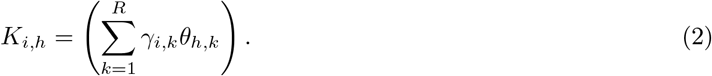

This gives a single value that takes both a species multidimensional preferences (i.e. fundamental niche) as well as the multidimensional resource abundances presented by a specific habitat into account. The better the fit, the higher *K*_*i,h*_, giving a species population density (if not further influenced by interactions or dispersal). Values for *γ*_*i,k*_ are iteratively assigned from a uniform distribution. The preference for the first resource *γ*_*i*,1_ is drawn from *U* (0, 1). The second is drawn from *U* (0, 1 − *γ*_*i*,1_) and the third receives the remainder (1-(*γ*_*i*,1_+*γ*_*i*,2_)). As the sum of *γ*_*i,k*_ is fixed at 1, a specialization for one environment comes at a cost for other preferences, see (Fig. 1B). We evaluate the dissimilarity of species pairs by evaluating the euclidean distance between environmental preferences of theses pairs (*E*_*ij*_, with similarity corresponding respectively to −*E*_*ij*_). We implement a minimum viable carrying capacity threshold, by setting growth rates *r*_*i,h*_ Eq.(1) of those species, where *K*_*i,h*_s are below 0.01 to 0. Otherwise *r*_*i,h*_ = 1.

##### Dispersal

Species dispersal is modelled by a diffusion-like exchange of populations between habitats. The exchange term *δ*_*i,h*_ (in Eq.1) is impacted be the net excess or deficit of a species population density compared with connected habitats. This is scaled by a dispersal coefficient D influencing the extent of abundance equilibration.

##### Simulations and sampling

The integration of population dynamics is performed with a Runge-Kutta Dormand–Prince (DOPRI) method (using scipy.integrate). Population dynamics and dispersal are intercalated with narrow time steps (0.02). Total simulation time is set so the vast majority of runs reach a steady state in community composition. All species abundances *N*_*i*_ *<* 0.01 are set to 0 post simulation (minimum viable carrying capacity). Post simulation we randomly sample 25 habitats and assessing species abundances. In later analyses we sample the habitat network space by volume. For this, we divide it into cubes with side lengths *l*. We trim the amount of habitats summed in one composite sample to a fixed amount. Species abundances from all habitats within a given volume are then summed. We discuss how varying numbers of habitats between samples in the Supporting information.

##### Determining co-occurrence

To determine a co-occurrence between two species we apply the most used but also naive approach (Fig. 1 F). The following criteria must be met: 1. A Spearman correlation coefficient |*ρ*| *>* 0.7 (the sign of *ρ* specifies positive or negative co-occurrence). 2. Correlation significance was tested by randomly reshuffling abundances within habitats (using *n* = 1000 permutations and significance threshold *p*_*cor*_ *<* 0.001, which is *p <* 0.05 Bonferroni corrected for *n* = 45 pairs).

## 3 Results

### 3.1 Imprints of drivers and their interplay in disconnected habitats

We start our analyses with our base scenario s1, in which we simulate community assembly in disconnected habitats with random resource compositions. Detailed information on scenarios referenced in this study can be found in the Supporting information (Tab. S2). Two factors drive community assembly: species environmental preferences and interactions. We first investigate how a species pairs (*i* and *j*) interaction coefficient *α*_*ij*_ and environmental preference similarity *E*_*ij*_ influence their co-occurrence. In Fig. 2 A positive co-occurrences are shown in red, negative ones in blue. We find a general pattern in which species pairs with larger |*α*_*ij*_| are correctly detected. For smaller coefficients, the value of *E*_*ij*_ determines the sign of co-occurrences. This can lead to positive co-occurrence between weakly competing pairs.

**Figure 2.**
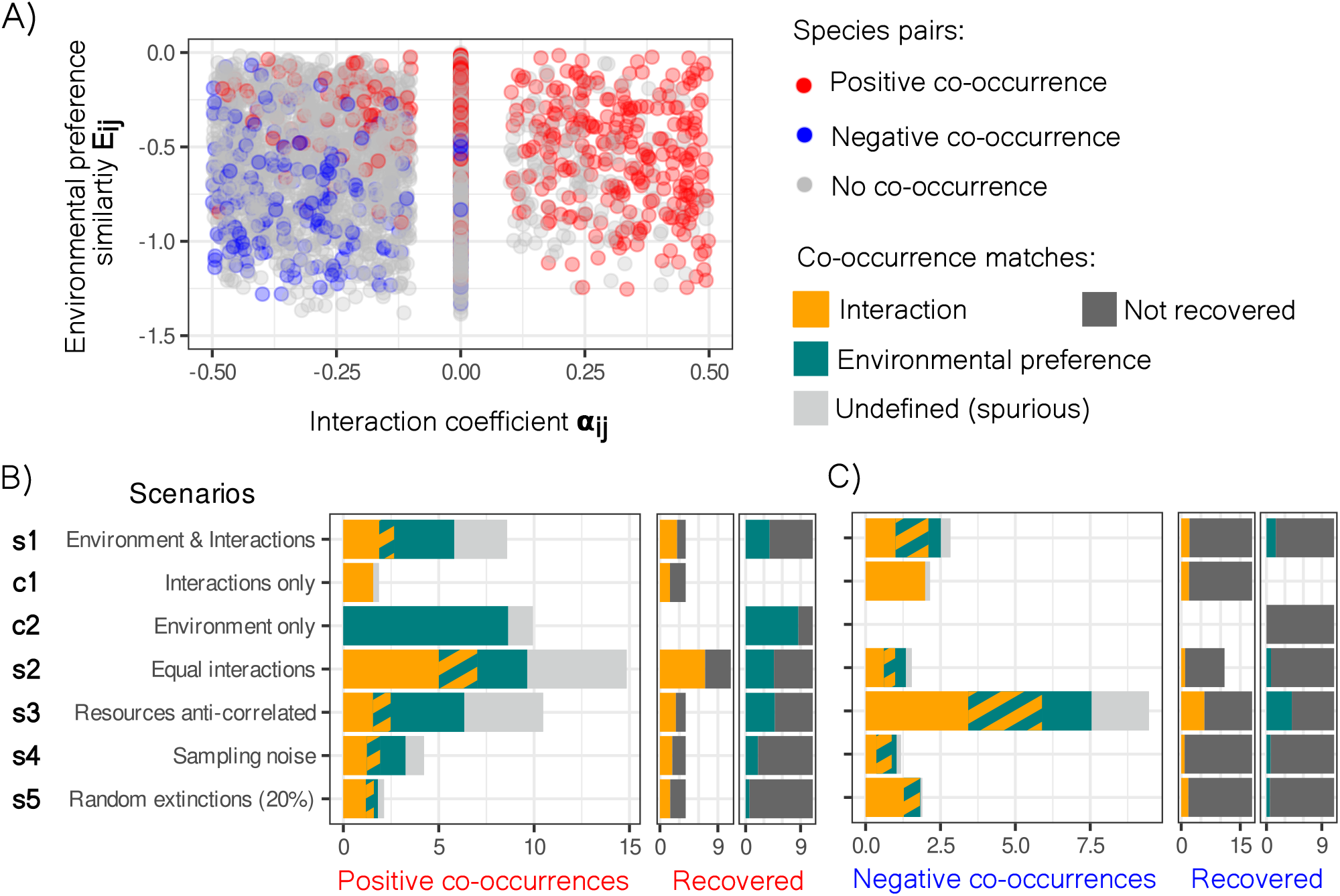
Imprints of interactions and environmental preference similarity on co-occurrence. A) In the depicted scenario interactions and environmental dynamics shape community composition (s1). All species pairs from 100 simulations are characterized by their respective interaction coefficient *α*_*ij*_ and environmental preference similarity *E*_*ij*_. Pairs are coloured by their co-occurrence: positive (red), negative (blue) or no co-occurrence (grey). B) Positive and C) negative co-occurrences are investigated for their possible origins in various scenarios. In the larger left panels we show the number of co-occurrences that match either the input interactions (orange), (dis-)similar resource preference (green), both (hatched) or spurious co-occurrences, which could not be traced to either (light grey). Of the two smaller panels next to this, the left shows how many of the input positive or negative interactions were actually detected by co-occurrences (orange) and how many were missed (dark grey). The right shows how many pairs with similar or dissimilar preference were detected (green) and how many were missed (dark grey). All values are averages of 100 runs.

Next, using our base scenario (s1), we quantify the number of co-occurrences that match our known input drivers (Fig. 2 B, C). To contextualise our findings we use additional scenarios, in which we vary the action of our two driving factors. First we look at positive co-occurrences. Acting independently (Fig. 2 B left; control scenarios c1 and c2) drivers are reconstructed with high precision (ca. 90% of co-occurrences matching either driver). Close to 50% of all input interactions were recovered (see B middle), while ~ 90% of all pairs with similar environmental preferences were recovered (see B right). When drivers are combined however (s1), the signal of interactions remains robust, but we observe a significant loss in the signal of environmental preference and a substantial number (~ 30%) of co-occurrences of spurious origin.^1^ This loss in the signal of environmental preference is even more pronounced when the number of positive interactions is high (s2). Next, we look at negative co-occurrences (Fig. 2 C). Here, the precision - so the percentage of co-occurrences that match a specific driver - in our base scenario (s1) for reconstructing negative interactions is high (~ 80%) compared to the precision for positive interactions (~ 25%; Fig. 2 C left). However, in this comparison we also see that the proportion of simulated negative interactions that could actually be recovered by co-occurrences is much lower (Fig. 2 C left and middle). This higher level in recovery for positive over negative interactions is highlighted when both interaction types are equally prevalent (s2). Compared to positive co-occurrences, negative co-occurrences are barely effected by species dissimilar resource preference, except in scenario s3. Here two resources are anti-correlated. Results closely resemble the pattern seen for positive co-occurrence in scenario s1. These results show that environmental dynamics only produce negative co-occurrences, when environmental factors are inversely distributed. Finally, we investigate the impact of stochastic effects such as sampling noise (s4) or random species extinctions (s5) on positive and negative co-occurrences. Here, the detection of similar environmental preference becomes obscured, while interactions remain more robustly detected.

In summary, positive interactions appear to be more readily detectable than negative ones. Nevertheless, we show that co-occurrences robustly reconstruct both positive and negative interactions even when environmental dynamics, or stochastic influences are considered. However, even in our simplified model we find that co-occurrences contain complex signals of both environment and interactions and that a significant portion can be entirely spurious.

### 3.2 Spatial structure and dispersal

Next we want to understand how the homogenization of microbial communities between different habitats affects observed co-occurrences. We establish a connectivity radius parameter *d*_*e*_, which determines the physical distance between habitats within which populations disperse. As expected, communities become more similar to each other with dispersal through increasing *d*_*e*_ (Fig. 3 B). The response of species richness to *d*_*e*_ is however non trivial (Fig. 3 A). All types of resource distributions display an initial increase and subsequent decrease (Fig. 3 A). Note, that while initially richness differs between these resource distribution scenarios, they converge to similar values at large *d*_*e*_. Interestingly, richness peaks are located around the point of network percolation (Fig. S2) and are slightly right-shifted with increased spatial structuring (i.e. resource patch size). With the increase in community similarity with increased *d*_*e*_, we also observe more of positive co-occurrences, most of which reflect shared environmental preference reducing the precision of inferring interactions (Fig. 3 C). However note, the presence of noise fully obscures this signal of positive co-occurrences at high *d*_*e*_, due to increased homogenisation (Fig. S3). We also see an initial increase and a rapid decline in the number of negative co-occurrences with *d*_*e*_, around the point of network percolation (Fig. 3 D). This can be explained by a loss in the detection of negative interactions, the primary driver of negative co-occurrence in our model (see previous section). What we see, results from high homogenisation via dispersal, where competitive species reach sufficient abundance across all habitats and an exclusion of their less competitive counterparts (see the decline in richness). Note, while we only depict co-occurrences for the scenario with mixed resource distributions (rs3) co-occurrence patterns are consistent across scenarios (Fig. S4).

**Figure 3.**
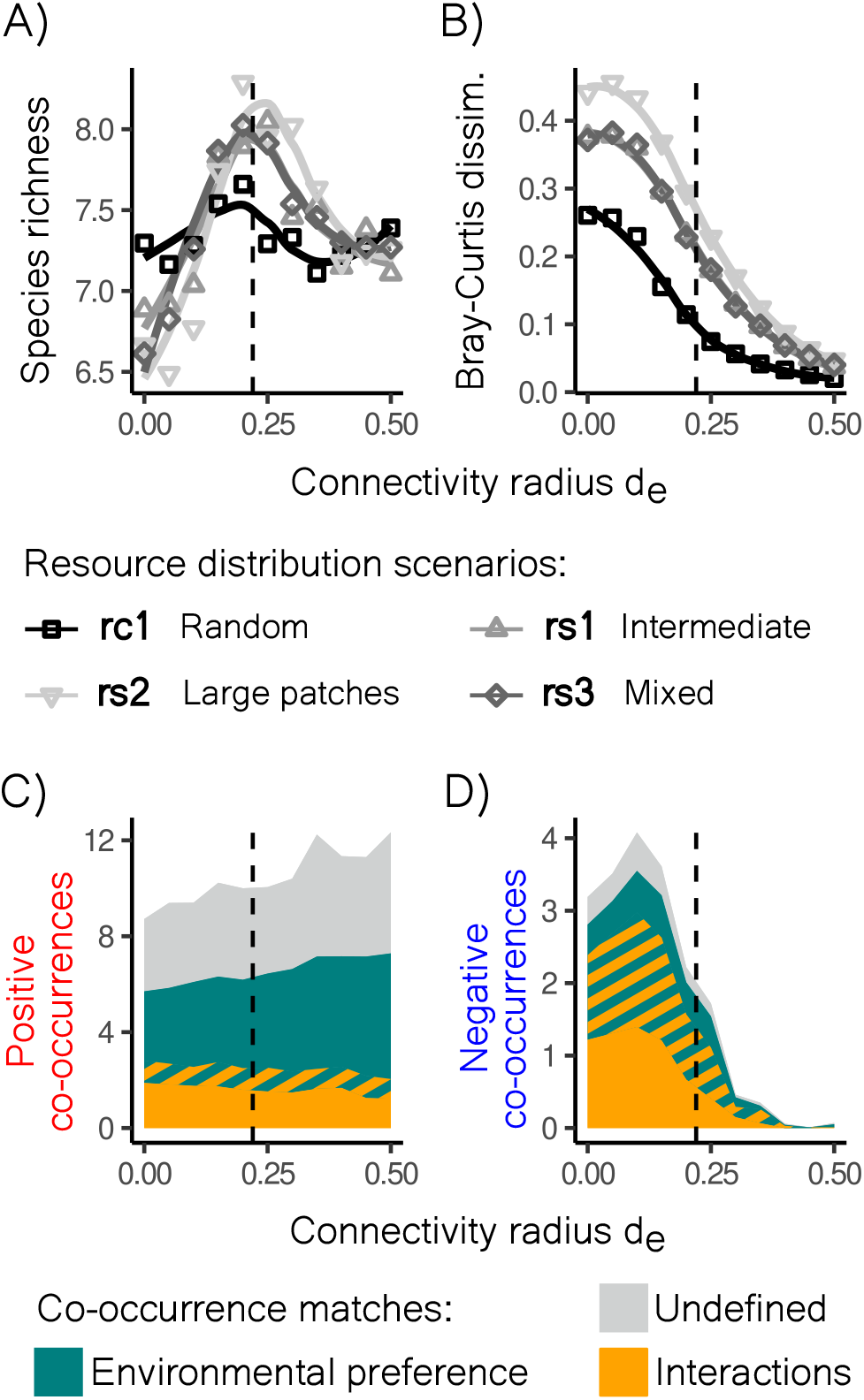
The effect of spatial resource distribution and dispersal on diversity and co-occurrence. The connectivity radius *d*_*e*_ controls the distance between which two habitats are connected and thereby enable the dispersal of populations. Dashed vertical lines show the point of percolation (*d*_*e*_ = 0.22), that is the distance at which habitat networks become fully connected. Responses of A) species richness and B) Bray-Curtis dissimilarity to *d*_*e*_ for various resource distribution scenarios. In the complete spatial randomness (rc1), intermediate (rs1) and large patch size (rs2) scenarios all three simulated resources are generated with equal spatial distribution inputs. For the mixed scenario (rs3) one resource is random, one intermediate and one has large patch sizes. C and D) show the number matching drivers for positive and negative co-occurrences respectively. Data is shown for the resource distribution scenario rs3. All values are averages of 100 runs.

### 3.3 Sampling scales

Studies based on sequencing data require material vastly surpassing what can be assumed individual micro-habitats. To understand how the scale sample units impact what we can learn from co-occurrence data, we sample at increasing volumes, sampling a fixed number of habitats and summing species abundances within (see Methods). We observe a general pattern independent of resource distributions: There is an increase in both positive and negative co-occurrences with sample volume (Fig. 4 B). We find that the more homogeneous the system from the start, the less the aggregation process inflates the number of co-occurrences detected. For the random scenario without spatial structure (rc1), we see almost no response to increased volume (Fig. 4 B). For scenarios in which all resources are spatially structured (rs1, rs2) however, we see an increase in co-occurrences. Here the precision for detecting interactions drops, but precision for environmental preferences remains unaffected (Fig. 4 C, D). This means that given sufficient spatial structuring of resources, for larger volumes the signal of environment becomes more evident. We see an intriguing case offered by scenario rc3, in which resources show varying levels of structuring. While dissimilarity is mostly equal to the intermediate distribution (rs1), it shows the highest increase in the number of co-occurrences with aggregation. While the drop in precision for detecting interactions mirrors the other spatially structured scenarios, precision for detecting environmental preference similarity drops substantially (Fig. 4 C, D). This likely stems from the challenge of detecting preferences for the randomly distributed resource (Fig. S5). In previous scenarios species abundance was primarily linked to the abundance of the resource they were most specialised for. Now the random resource is found in near equal quantities across all samples, while the spatially structured resources maintain substantial variation. So where before it was three resources determining species abundance patterns, it is now mainly two. Consequently, the abundance of more species is found fluctuating with fewer resources leading to more co-occurrences. As this includes those dominantly preferential for the random resource, fewer resulting co-occurrences can be traced to close similarity in environmental preference.

**Figure 4.**
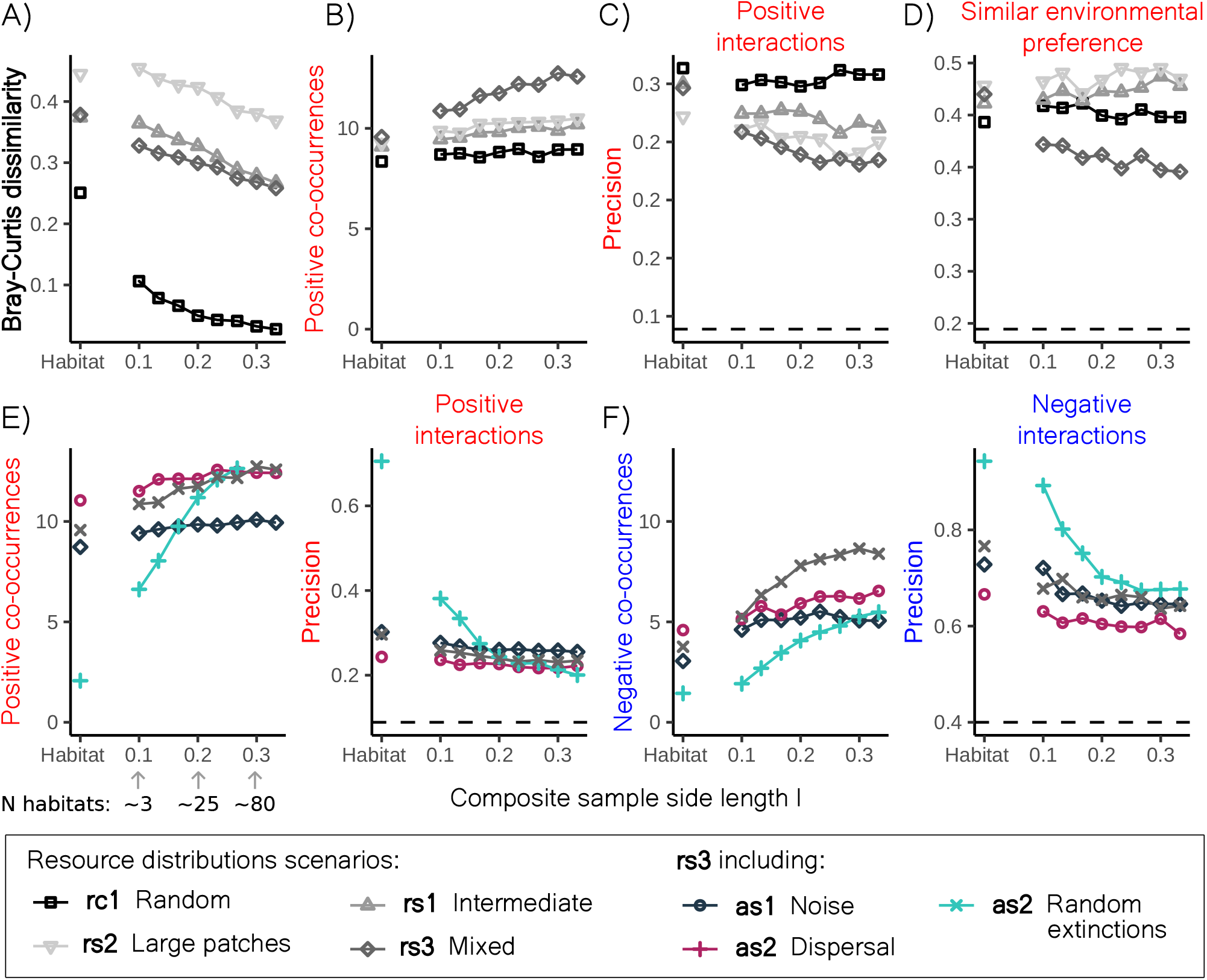
Co-occurrences and the imprint of drivers at increasing sampling volumes. Our analysis first focusses on how results are constrained by the spatial distribution of resources (scenarios found in Fig. 3). A) The mean decrease in dissimilarity between samples, B) mean number of observed positive co-occurrences and the corresponding precision for detecting C) positive interactions and D) environmental preference similarity are depicted as a function of the number of habitats contained within composite samples. Next, by modifying the mixed resource distribution scenario (rs3) we analyse how confounding influences from sampling noise (as1), dispersal (as2) and random extinctions (as3) further impact our results. E) and F) show the number of positive and negative co-occurrences (left) and the precision for detecting positive or negative interactions (right) for these scenarios respectively. Dashed horizontal lines in C), D), E) and F) show values for random precision. All values are averages of 100 runs.

To include further dynamics encountered in microbial studies, we introduce additional scenarios, all of which are based on the resource distributions of rs3. The co-occurrence signal at larger volumes is generally robust to the effect of sampling noise (as1) in spatially structured scenarios (Fig. 4 C, D). Even when noise is large only decreases in the number of observed co-occurrences with scale are only moderate. Note, when resources are extremely heterogeneous however (i.e. random distribution scenario) sampling noise drastically reduces the the number of co-occurrences (Fig. S6). In contrast to noise, stochastic extinctions (as3) heavily deter observation of co-occurrences at small volumes. Unlike sampling noise, the effects of random species extinctions seem to average out when sampling larger volumes, especially for positive co-occurrences, where results are indistinguishable from those of the base scenario rs3. Finally, it is important to mention why we control for an equal numbers of habitats in composite samples. If not, spurious positive co-occurrences arise purely from variation in the number of habitats, which heavily skew the abundance of species in composite measurements (Fig. S7).

## 4 Discussion

### 4.1 Main findings

In this study, we developed a spatially and environmentally explicit generalized Lotka-Volterra (gLV) model including meta-community dynamics. Our goal was to investigate how various dynamics and properties of microbial ecosystems affect species co-occurrence. Our findings present a nuanced perspective. On one hand, we show that the signal of strong species interactions can still be detected despite environmental confounders, with positive interactions generally being easier to detect than the negative ones. On the other hand, we show that environmental confounders introduce additional complexity, not only by making it challenging to disentangle the underlying drivers but also by generating a considerable fraction (approximately one-third in our case) of co-occurrences that arise from unknown indirect or spurious effects. We also explored the effects of homogenization on observed co-occurrence patterns and the reconstruction of community assembly mechanisms. Two forms of homogenization were considered: intrinsic (resulting from dispersal between meta-communities) and extrinsic (stemming from sampling processes, such as aggregating large volumes of material for sequencing). Within the parameters analysed, both forms inflate positive co-occurrences. This inflation results from enhanced detection of environmental drivers, simultaneously reducing the precision for reconstructing interactions. For negative co-occurrences however, we see opposite responses, meaning an inflation with extrinsic-and a reduction with intrinsic homogenisation. Note, both processes fundamentally differ, as one (intrinsic) affects population dynamics and can induce global extinctions, while the other (extrinsic) maintains more variation in community similarity at large volumes when resources are spatially distributed. Our findings highlight the critical need to consider both microbial community composition and environmental inter-sample variability to accurately identify the drivers of community assembly with the use of co-occurrence analysis, see Table. 4.1 for summary.

**Table 1:** Summary of the effect of two types of homogenization ((i) intrinsic, driven by dispersal and connectivity between microhabitats; (ii) extrinsic, realized by aggregating and mixing several habitats into a single sample) on co-occurrences observed. ^*^ depends on the resource distribution and scenarios considered (presence of noise).

|  | positive |  | negative |  |
| --- | --- | --- | --- | --- |
| homogenization | (i) dispersal | (ii) sampling | (i) dispersal | (ii) sampling |
| co-occurrences | ↑ | ↑ | ↓ | ↑ |
| precision of interactions | ↓ | ↓ | — | ↓ |
| precision of environment | ↑ | ↑* | — | ↓ |

### 4.2 Implications for co-occurrence network analysis

Numerous studies have advised caution using co-occurrence patterns to infer microbial interaction webs (Blanchet et al., 2020; Guseva et al., 2022; Carr et al., 2019). Studies leverage numerically generated data with known parameters to investigate their ability to reconstruct interactions (Berry et al., 2014; Weiss et al., 2016; Pinto et al., 2022). However, despite these advancements, a significant gap remained: Approaches have mostly overlooked environmental dynamics as explicit drivers of co-occurrence patterns. Furthermore, little attention has been given to understanding how spatial context, such as the heterogeneity in environmental factors and the connectivity of habitats might influence co-occurrence signals. This oversight heavily limits our ability to interpret what information species co-occurrences themself and consequently co-occurrence network structures as a whole might hold. Addressing this complexity is imperative to improve current (often highly speculative) interpretations. Our analysis so far has focussed on analysing co-occurrences ability to reconstruct drivers independently. Here we want to illustrate how our main drivers shape network structure with networks that depict positive co-occurrences driven by: i) environmental factors (Fig. 5 A); ii) interactions (Fig. 5 C) and iii) a combination of both drivers (Fig. 5 B). In i) one can see that generalist species are found central and modules contain environmental specialists. While in ii) the position of species in networks can reflect its role in the community (Berry et al., 2014). When both drivers interplay iii) we see a drastic reorganisation of network structures. Modules become amalgamations of fewer specialists and their interaction partners, meaning both drivers interact to produce observed structures. Within modules we see how many indirect dependencies produce unaccounted for co-occurrences (black edges). We also see that the inclusion of environmental factors can only give limited context. While we see a module with three specialists correctly associated with the cyan resource, we also how two specialists for the pink resource that are associated a false resource influenced by interactions. This highlights the fact that identification of environmentally driven co-occurrences through methods such as sign pattern analysis are best approached with caution (Deutschmann et al., 2021; Lima-Mendez et al., 2015). While species-environmental factor associations can provide context when imposed on co-occurrence networks, information such as trait or metabolic complementarity assessments would provide much needed additional context suitable for implicating origins of co-occurrence (Zafeiropoulos et al., 2024).

**Figure 5.**
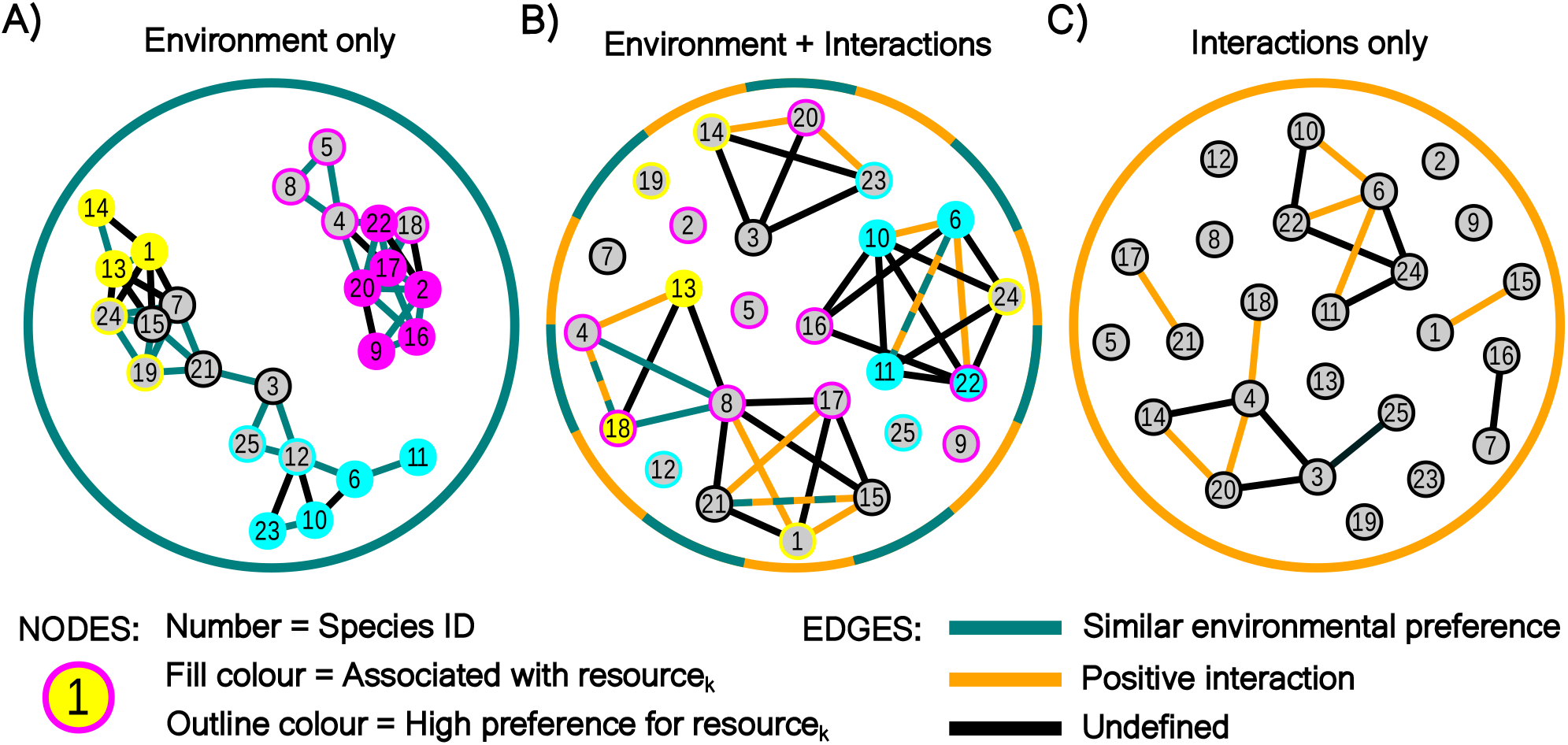
Exemplary networks for simulations of a ‘simple system’. Networks are constructed from data simulated with the main settings of scenarios A) c2, B) s1 and C) c1. The only difference is that here we simulate 25 species. Of all 300 pairs 10% engage in positive (= 30), 40% in negative interactions (= 120) and 50% do not interact. The upper and lower 25% of environmental preference similarities (*E*_*ij*_) are detected similar or dissimilar (= 75 each). For these three simulations inputs are mostly the same (interaction matrix, preferences, resource abundances, etc.). The control scenarios effectively deactivate the other drivers as mentioned in the first result section. The figure is meant to illustrate how networks imprinted by either driver how topology changes when drivers are combined.

### 4.3 Models for analysing co-occurrence

In this work, we chose to utilize gLVs to simulate the assembly of interacting communities. However, we recognize that analysing co-occurrence on a broader range of numerically generated data would provide valuable insights. Comparative approaches, integrating alternative modelling frameworks, offer significant potential for advancing our understanding of the limitations inherent to various inference methods, while also revealing possible biases or constraints unique to specific models. The question of which framework is most suitable for describing microbiome dynamics remains a topic of active debate in the field (Picot et al., 2023; Berg et al., 2022). Beyond gLVs, other commonly used models include MacArthur’s consumer-resource framework, individual-based models and game-theoretic approaches, each capturing distinct aspects of microbial dynamics and offering varying levels of detail and complexity. A critical point of discussion involves the relevance of higher order interactions among taxa and their effect on the community structure (Ishizawa et al., 2024; Cui et al., 2024; Sidhom et al., 2024; Gibbs et al., 2022). However, as our study primarily focussed on the influence of environments on species co-occurrences, we limited our analysis to the simplest and arguably the most popular framework. We also note that in this context, we have chosen to work with a simple set of interaction parameters (randomly assigned and symmetric) to define our communities. Again, to focus our analysis on environmental dynamics we limited our range of tested interaction coefficients. Our goal was for coefficients to broadly represent prevalence of positive and negative interactions from empirical observations (Kehe et al., 2021; Palmer et al., 2022) and to select interaction strengths that would maintain co-existence of competing species pairs (further discussed in Supporting information: Interaction coefficients). In this work, we have not explored the effect of possible interaction network topologies on co-occurrence recovery in depth. In this context we recommend Pinto et al., 2022 for a comprehensive overview on how various forms of interactions imprint on co-occurrences (in absence of the environmental factors) and an extensive discussion on the utility of gLV simulated data for such studies in general.

### 4.4 Environmentally explicit dynamics

We highlight that our model allows for an ‘environmentally explicit’, consistent response across habitats, that allows for similar resource preference to coherently imprint on species co-abundances, within a framework that rests on the fundamental niche concept (Hutchinson, 1957; Colwell et al., 2009). This framework allows different species to respond to (i.e. prefer) different environmental factors, describing a gradient of specialists and generalists. We note that in our implementation, environmental factors confer linear, quantitative benefits to species abundance. Therefore in our context we refer to these factors as ‘resources’. However, in reality microbial responses to environmental variables can be non-linear, or show growth optima for parameters such as temperature, oxygen concentration, and pH (Yu et al., 2025; Lee et al., 2024; Lin et al., 2024). Here scenario s3 (Fig. 2 B, C) is noteworthy, where we inversely correlate the distribution of two resources we also imitate possible opposing preferences for single environmental factors (think oxic vs. anoxic bacteria). In addition, for simplicity, our implementation assigned preferences and interactions fully independently of each other. While this simplification offers a starting point for analysis, it is important to note that these factors are often closely linked in natural systems. For example, negative interactions might arises from overlapping ecological niches. Finally, in many natural systems there can be a less clear separation between population dynamics (driven by interactions) and environmental filtering dynamics, resulting in tight species-environment feedbacks (this can be the case of pH) (Ratzke et al., 2018). Such feedbacks can have a non-trivial impact on the spatial structure of the community, as can be exemplified by the system from Liautaud, Nes, et al., 2019. In this theoretical study sharply delineated patches containing consistent community compositions were shown to emerge along environmental gradients. However their boundaries can be blurred by dispersal or straightened by such species-environmental feedbacks (Liautaud, Barbier, et al., 2020). This would have a strong effect on the distribution of species abundance data and therefore their correlations.

### 4.5 Meta-community dynamics

Our work also contributes to the growing body of research emphasizing the importance of understanding how microbial taxa are distributed across spatially connected microhabitats. Previous theoretical studies based on gLVs, as well as on individual-based models, highlight the intricate relationship between dispersal and biodiversity (Denk et al., 2022; Gravel et al., 2016; Mattei et al., 2024; Bickel et al., 2020; Garcia Lorenzana et al., 2024). While they mostly concentrate on interactions, demographic fluctuations and habitat connectivity, our work is the first to show how the physical distance of dispersal and the spatial distribution of environmental factors interplay. We see that the spatial extent of environmentally similar patches is directly related to the distance dispersal must cover to produce biodiversity optima. In addition, we demonstrate how this meta-community framework can be used to investigate how underlying dynamics are imprinted on co-occurrence patterns, offering new insights for empirical studies.

### 4.6 Simulating holistic community assembly

Our model simulates community data across connected habitats, embedded in heterogeneous environment landscapes. Our model offers a holistic approach, where communities emerge from the interplay of fundamental ecological processes with tunable spatio-environmental system characteristics, going beyond other models that cover certain aspects of community assembly (Ruffley et al., 2019; Plas et al., 2015; Etherington et al., 2019; Fraboul et al., 2023; Lytle et al., 2023). As various tools try to quantify the effects of different assembly drivers (Borcard et al., 1992; Schulz et al., 2025; Guisan et al., 2000; Ovaskainen et al., 2017; Warton et al., 2015), it could be of broad interest for generating ecological test data, benchmarking various analytical tools, and investigating their inference potential. An emerging approach in the field of co-occurrence network analysis known as “coupling networks” focusses more heavily on the integration of environmental variables (Ochoa-Hueso et al., 2021), either via partial correlations (Kurtz et al., 2015) or even via model-based methods (Rao et al., 2021). To evaluate how these tools perform, so how f.ex. the structure of coupling networks relate to system resilience, we propose applying our model framework. Empirical ecological data is inherently complex and our model provides a more realistic representation of natural systems dynamics. This level of intermediate complexity is required for bridging theoretical insights with practical observations.

### 4.7 Conclusion

In this work we propose a new framework for modelling microbial systems in the context of environmental heterogeneity, use it to produce co-occurrence patterns and show how community assembly drivers interact producing complex imprints on co-occurrences. Our results demonstrate that detecting true interaction signals from co-occurrence data is substantially impeded by both extrinsic (e.g., intentional mixing of heterogeneous samples, such as done for systems as soil) and intrinsic (e.g., such as probably happens naturally in well-mixed marine environments) homogenization. Note that in both contexts, homogenization masks ecological signals, complicating inference of species interactions and niche preferences. We conclude by given some recommendation for practical applications of co-occurrence analysis. We recommend to (1) characterize micro-scale spatial variation of environmental factors; (2) optimize sampling strategy to match environmental heterogeneity; (3) interpret positive co-occurrences with caution. Finally we also strongly recommend for researchers to use numerical datasets to understand the limits of co-occurrence analysis.

## Acknowledgements

SD, CK and KG have received funding from the European Research Council (ERC) under the European Union’s Horizon 2020 research and innovation programme (consolidator grant agreement No. 819446, granted to CK)

## Footnotes

1 Co-occurrences with undefined origin are referred to as spurious due to their limited interpretative value. This does not mean they are non-deterministic. They may arise from indirect effects of interactions or from very weak similarity in environmental preference.

