## Supplementary Material for "Environmental context shapes microbial co-occurrence"

August 5, 2025

**Abstract**

**S2 Interaction coefficients $\alpha_{ij}$** **6**
**S3 Scenario descriptions** **7**
**S4 Extended Results and Discussion** **8**
S4.1 Sampling scales . . . 8

### S1 Extended Material and methods

To make our study more accessible to a general audience from a range of disciplines in ecology, we decided to keep the Material and methods section in the main manuscript short an concise. In this section we give a more detail oriented overview of the methods used in this study.

#### S1.1 Table of model variables

For an easier overview and referencing we provide a table of the variables and indices used in this study S1.1.

| Variable [index] | Description |
|---|---|
| $L^3$ | Analysed spatial volume of resources and habitats ($L = 1$) |
| $H$ [$h$] | Number of habitats (indices if multiple = [$h1$,$h2$,...]) |
| $d_e$ | Radius within edges are set between habitats (in scale of $L$) |
| $R$ [$k$] | Number of resources (indices if multiple = [$k1$,$k2$,...]) |
| $\lambda_k$ | Exponent of the power law spectrum for resource distribution (= landscape) |
| $\phi_k(x, y, z)$ | Resource abundance within a landscape voxel |
| $\theta_{k,h}$ | Resource abundance in a habitat |
| $\Theta_{k,h}$ | Sum of resource abundances |
| $S$ [$i$] | Number of species (index for pair = [$ij$]) |
| $\gamma_{i,k}$ | Species resource preference |
| $K_{i,h}$ | Species habitat carrying capacity |
| $r_{i,h}$ | Species habitat growth rate |
| $\alpha_{ij}$ | Species inter-specific interaction coefficient |
| $D$ | Dispersal coefficient |
| $N_{i,h}$ | Species habitat abundance |
| $l$ | Composite sample side length (in scale of $L$) |

Table S1: List of variables and indices used in our model, with a short description.

#### S1.2 A network of habitats in spatially explicit environments

We consider an observation volume of dimensions $L^3$ ($L = 1$). Within this volume we construct a random geometric graph (undirected and without self-loops), characterized by a given number of nodes-habitats $H$ and edges between habitats allowing species to disperse. The number of habitats is set to $H = 25$ in the initial analyses. $H = 300$ when we move to the spatially explicit system with dispersal. Finally, $H = 3000$ when sampling at increasing volumes. Habitats are simulated with random positions in the observation volume. The average connectivity of these habitats $\langle c \rangle_h$ is varied by adjusting the parameter $d_e$ (fraction of $L$). Smaller $d_e$ implies fewer connections and $d_e > 1.73L$ describes a fully connected network. The adjacency matrix that represents this network of habitats is denoted by $W$, and $W_{h1,h2}$ assumes value 1 in the presence and 0 in the absence of an edge. Each habitat is characterized by a location ($x_h$,$y_h$,$z_h$), given as fractions of $L$ and a certain number of edges that connect it to other habitats. A habitats resource composition $\theta_{k,h}$ is informed by its location, which is projected into three different resource landscapes ($R = 3$). These are modelled as scale-free Gaussian Random Fields in $100^3$ voxel grids, which are defined by a power law spectrum with exponent $\lambda$ (Cadiou, 2022). This form of spectra defines a resource landscape with resource abundance values in every voxel $\phi_k(x, y, z)$ characterized by a variety of different spatial structures. Given the coordinates ($x_h$,$y_h$,$z_h$) of each habitat $h$ in space, we assign the abundance of resource $k$ in it as $\theta_{k,h}$ with the punctual abundance of the resource in space $\theta_{k,h} = \phi_k(x_h, y_h, z_h)$. $\lambda$ gives the strength of spatial structuring as it defines the prevalence of small scale structures, we select three representative settings: (i) $\lambda = 0$ gives complete spatial randomness; (ii) intermediate $\lambda$ field is dominated by high frequency fluctuations ($\lambda = 5$); (iii) large $\lambda$ can be described by large patch structures ($\lambda = 10$). In total we test five different resource distribution scenarios. We assume that species have different responses to different resources and generate three resource landscapes independently per run ($R = 3$). The spatial heterogeneity of each scenario therefore results from a set of distributions $\lambda$; one for every resource. For the first three scenarios $\lambda$ is always the same. Our control with no spatial structuring (i.e. complete spatial randomness) is defined **rc1** $= \{\lambda_0, \lambda_0, \lambda_0\}$. This scenario is used in our results of a 'simple system' in Fig. 2 (apart from scenario **s5**). The intermediate spatial structuring scenario is defined **rs1** $= \{\lambda_5, \lambda_5, \lambda_5\}$. The scenario with maximum spatial structuring (i.e. large patch sizes) is defined **rs2** $= \{\lambda_{10}, \lambda_{10}, \lambda_{10}\}$. These allow us to analyse three naive scenarios of different spatial heterogeneity (i) fully random; (ii) fractal structures and (iii) patchy. We include the fourth scenario **rs3** $= \{\lambda_0, \lambda_5, \lambda_{10}\}$ to give a possibly more realistic scenario where distribution patterns of resources vary. All of these scenarios are used in the results of Fig. 3 and Fig. 4. In a final scenario we investigate how an inverse distribution of resources might effect our observations. Here the first and second resource $k1$ and $k2$ are inversely distributed in gradients. A gradient from 1 to 0 (squared values) runs from top to bottom for $k1$ and from bottom to top for $k2$. $k3$ is then randomly distributed ($\lambda_0$). Visualisations of example resource landscapes can be found in Fig. S1.

#### S1.3 Simulation of population dynamics

**Dispersal** The time evolution of the biomass modelled in Eq.(**??**) is complemented by a diffusion-like exchange of species abundances between habitats. The exchange term is given by:

$$\delta_{i,h1} = D \sum_{h2=1,h1\neq h2}^{H} W_{h1,h2}(N_{i,h1} - N_{i,h2}), \tag{1}$$

where D is the dispersal coefficient. The strength of D influences the extent of the equilibration of abundances, or homogenisation but is constrained by the connectivity of habitats. The number of connected habitats ($h1$,$h2$,...) found in the adjacency matrix $W$ varies with the settings for the connectivity radius $d_e$. The relationship between $d_e$ and overall network connectivity, relies on the number of habitats that are simulated. In Fig. S2 we show how mean node degree increases with $d_e$ for 300 (B) and 3000 (D) simulated habitats. While the shape of this relationship is not effected by the number of habitats, the value of $d_e$ at which the network percolation point is reached is impacted. Both the number of components (i.e. number of disconnected nodes or node clusters) and the average path length (average number of nodes that lie between two random nodes) can be used to approximate the percolation point. With $H = 300$ this point is reached at $d_e \sim .22\%$ (A), while it is already reached at $d_e \sim .11\%$ when $H = 3000$.

**Species environmental preference** One of the main goals of our model was to implement a simple but realistic fundamental niche dynamic. For this we simulate species with varying degrees of specialisation for a range of resources. We assume that specialization of a species for one resource can only come at a cost of decreasing specialization for another resource. This trade off is modelled using a constraint for $\gamma_{i,k}$ values, which is given by a constant sum, i.e. we assume $\sum_{k=1}^{R} \gamma_{i,k} = 1$. The values $\gamma_{i,k}$ are selected randomly from a uniform distribution $U(0, \max)$. For this we decide a random order of $k$'s, meaning in each step a

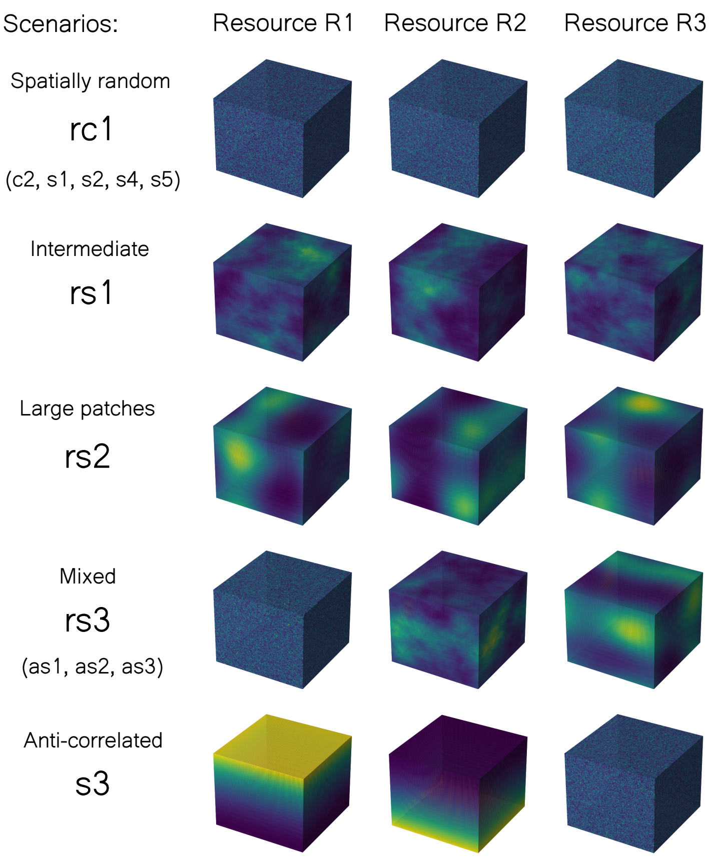

Figure S1: Examples for the spatial distributions of environmental factors we simulate are presented for each scenario analysed in the manuscript. What we call 'resource landscapes' are shown in columns for each resource we simulate ($R = 3$). Rows show the various scenarios referred to in the main manuscript. Brighter yellow colours represent higher resource abundance and darker blue colours represent low abundance.

random $R$ from $k1$, $k2$ or $k3$ is selected. If $k1$ is first, then $\gamma_{i,k1}$ is assigned a value from $U(0, 1)$. The next could be $k2$, for which now $\gamma_{i,k2}$ is sampled from $U(0, 1 - \gamma_{i,k1})$. The remaining $k3$ will now receive $\gamma_{i,k3} = 1 - (\gamma_{i,k1} + \gamma_{i,k2})$, which makes $\sum_k^R = 1$. Through this sequential assignment some species can have high resource specialisation if the first values selected are very high. Species resource preferences can be visualised in a multidimensional space and we term the position of a species in this space their environmental preference, which can be understood as their environmental niche. To analyse how species similarity in environmental preference might influence their co-occurrence we calculate the euclidean distance between species pairs

$$E_{i,j} = -\sqrt{\sum_{k=1}^{R} (\gamma_{i,k} - \gamma_{i,k})^2} \tag{2}$$

thereby receiving a metric, where low negative values represent high similarity in overall preference for resources. Higher negative values are produced when species resource preferences are dissimilar.

**Simulations and sampling** We intercalate steps of population dynamics and dispersal. As we set the step sizes for integration very small (0.02; 5000 steps for a total simulation time of 100) this allows us to keep a balance between the two separate dynamics during simulation. After simulation we randomly sample

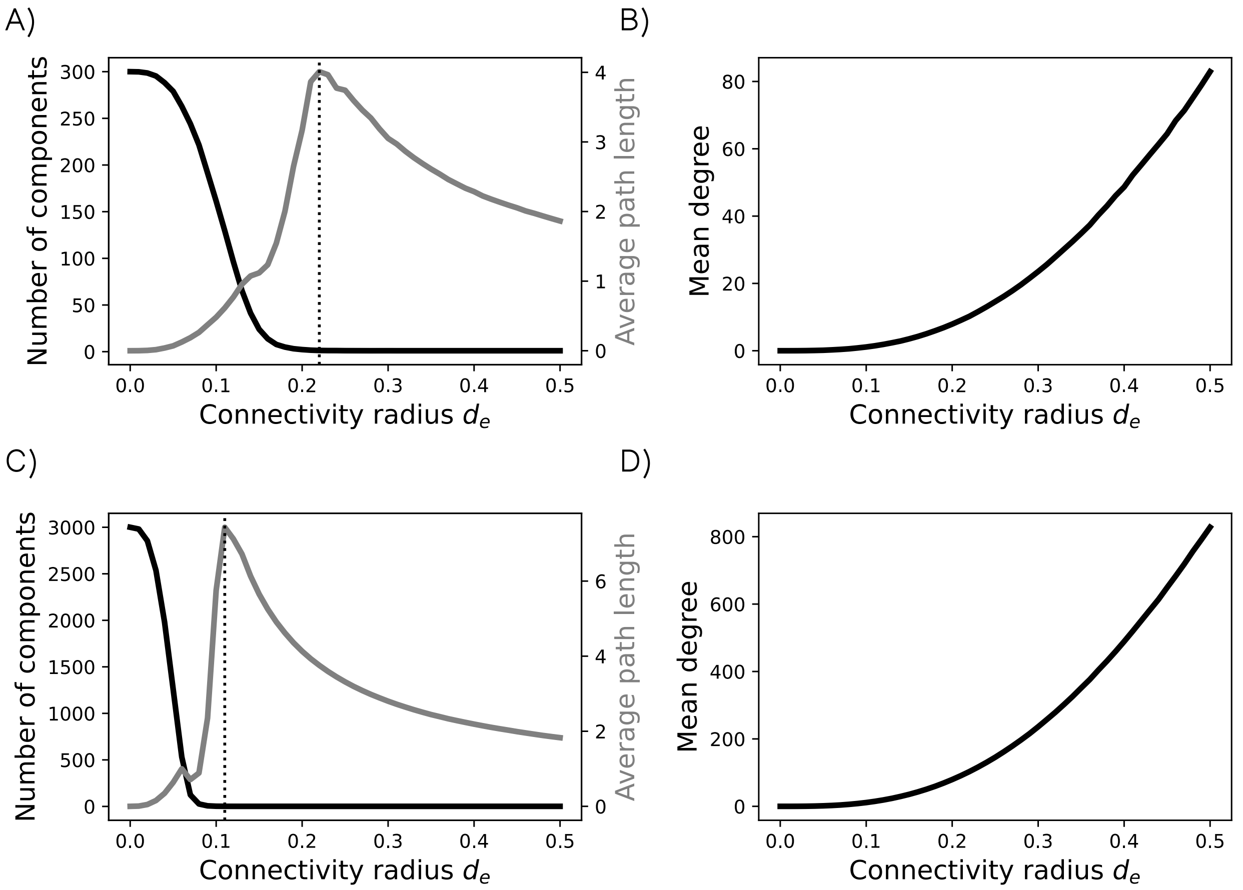

Figure S2: Network connectivity properties as a function of the connectivity radius $d_e$ and the number of simulated habitats. A and C show the number of components in a network as well as the average path length between nodes. B and D) show the mean node degree. A and B are based on simulations with $H = 300$, while C and D are based on simulations with $H = 3000$.

$G$ number of samples. In all analyses we fixed this number to $G = 25$, as preliminary analyses showed that increasing this number did not meaningfully change our results. This finding was robust for the parameters we selected but might vary for other settings. When analysing different volumes a sample is sourced from a certain volume $l^3$ (fraction of $L$). For this we divide total space $L^3$ into cubes with side lengths $l$ and sample cubes meaning no two samples cover the same space. Due to the random coordinates of habitats in the observation volume ($L^3$) sampled cubes can contain varying numbers of habitats. We decided to control for a fixed number of habitats contained within a sample cube before downstream analysis (why is explained in the section 'Simpsons paradox'). For this we rank cubes (given a specific side length $l$) by the number of habitats they contain. The $G = 25$ highest ranking cubes are sampled and randomly trimmed so all contain the same number of habitats, i.e. to the number of habitats in the cube ranked 25th. For further analysis species abundances of the remaining habitats are summed and the cube is treated as a single sample.

### S2 Interaction coefficients $\alpha_{ij}$

As the analysis of our study focuses primarily on the factors of community assembly related to environments and space, we limited the scope of interaction parameters tested. Rather than aiming for full empirical realism, our parameter choices represent a compromise: they are loosely inspired by patterns observed in natural systems but were ultimately selected for analytical tractability. In particular, we aimed to capture general trends—such as the high prevalence of non-interacting species, a dominance of competitive interactions, and a minority of facilitative ones (Palmer et al., 2022), all while deliberately excluding more complex or asymmetric types of interactions (e.g., amensalism, commensalism, and predation). This allowed us to explore assembly dynamics in a controlled, interpretable setting, while still being loosely grounded in empirical insights. Kehe et al., 2021 produced an extensive dataset analysing pairwise interactions between 20 bacterial strains in 40 different media (total of 180,408 pairs). To get an idea of not only the signs but also the interaction strengths, so the quantitive enhancement or reduction of a pairs abundances we analysed their primary data. Here we learned that 29% of interactions were deemed fully competitive (45% including amensalism). Species suppressed each others biomass by 45% (both mean and median). In our settings 40% of species engage in competition with an average suppression of 23% and a maximum of 33%. Next, 5% of all associations were found mutualistic (19% including commensalism). Here the mean enhancement of abundances was 276% (median 80%). In our case 9% of species engage in facilitation and the average enhancement was 43% and the maximum was 100%. Interaction strengths therefore tended to be weaker in our settings than in these data. Our goal was to allow the coexistence of competitive species and to hinder communities from being heavily dominated by mutualistic pairs. These data were however attained in artificial settings in pairwise incubation on simple substrates, meaning these results might not reflect pairwise interactions in natural systems. The simplified interaction settings analysed in our manuscript therefore strike a deliberate balance between empirical realism and analytical interpretability, allowing us to capture key ecological trends while avoiding the confounding complexity of more intricate interaction dynamics. We hope that future research will be able to expand on how more diverse forms of interactions impact our results.

### S3 Scenario descriptions

This section aims to give an easier overview of the scenarios in our analysis. Methods in the main manuscript as well as the Extended Material and methods above give an overview of the main settings applying to our base scenario **s1**. All following scenarios modify these settings in few particular aspects. Tab. S2 gives information on which settings were altered to produce the various scenarios in our analysis. The spatial resource distributions used for scenarios are described above and depicted in Fig. S1.

Table S2: All scenarios that are shown in the results are listed including a short description and the Figures they appear in.

| **Scenario** | **Description** | **Main figure** |
|---|---|---|
| **s1** | Base scenario (described in methods) | Fig. 2, Fig. 5 |
| **cs1** | Random $K_{i,h}$ from $N(0.27, 0.13)$. These values are taken from the distribution of $K_{i,h}$ values in s1 | Fig. 2 |
| **cs2** | All interaction coefficients $\alpha_{ij}$ are set to 0 | Fig. 2 |
| **s2** | Competition and mutualism are equally prevalent (both 11 pairs) | Fig. 2 |
| **s3** | Opposing environmental gradients for the first two resources are simulated, effectively leading to a strong anti-correlation in the abundance of these two resources across sampled habitats | Fig. 2 |
| **s4** | The effect of imprecise measurements is analysed by adding random noise to species abundances in habitat or composite samples. Random values from $N(0, \sigma_i \, \xi)$ are generated, where $\sigma_i$ is the standard deviation of a species' abundances across samples and $\xi$ scales the noise strength (here $\xi = 0.25$). | Fig. 2 |
| **s5** | Random extinctions are simulated by excluding a random subset of species when initially populating habitats. Here only 8 of 10 species are initialised in any given habitat. | Fig. 2 |
| **rc1** | Resources show spatially random distribution (all $\lambda = 0$) | Fig. 3, Fig. 2 |
| **rs1** | Resources show intermediate spatial distribution (all $\lambda = 5$) | Fig. 3, Fig. 4 |
| **rs2** | Resources show spatial distribution with large patches (all $\lambda = 10$) | Fig. 3, Fig. 4 |
| **rs3** | All three resources show different distributions: one is spatially random ($\lambda_{R1} = 0$), one intermediate ($\lambda_{R2} = 5$) and one large-patched ($\lambda_{R3} = 10$) | Fig. 3, Fig. 4 |
| **as1** | Modified rs3. Sampling noise added as in s4 with $\xi = 0.25$ | Fig. 4 |
| **as2** | Modified rs3. Dispersal in an intermediately connected habitat network: dispersal coefficient $D = 0.1$ and habitat connectivity radius $d_e = 0.1$. The relationship of $d_e$ and the percolation point is illustrated in Fig. 1. | Fig. 4 |
| **as3** | Modified rs3. 80% random extinctions as in s5 | Fig. 4 |

### S4 Extended Results and Discussion

Here we present the Supplementary figures referenced in the main manuscript. Figures are presented under their corresponding Result section titles and are arranged in the order in which they appear in the main manuscript.

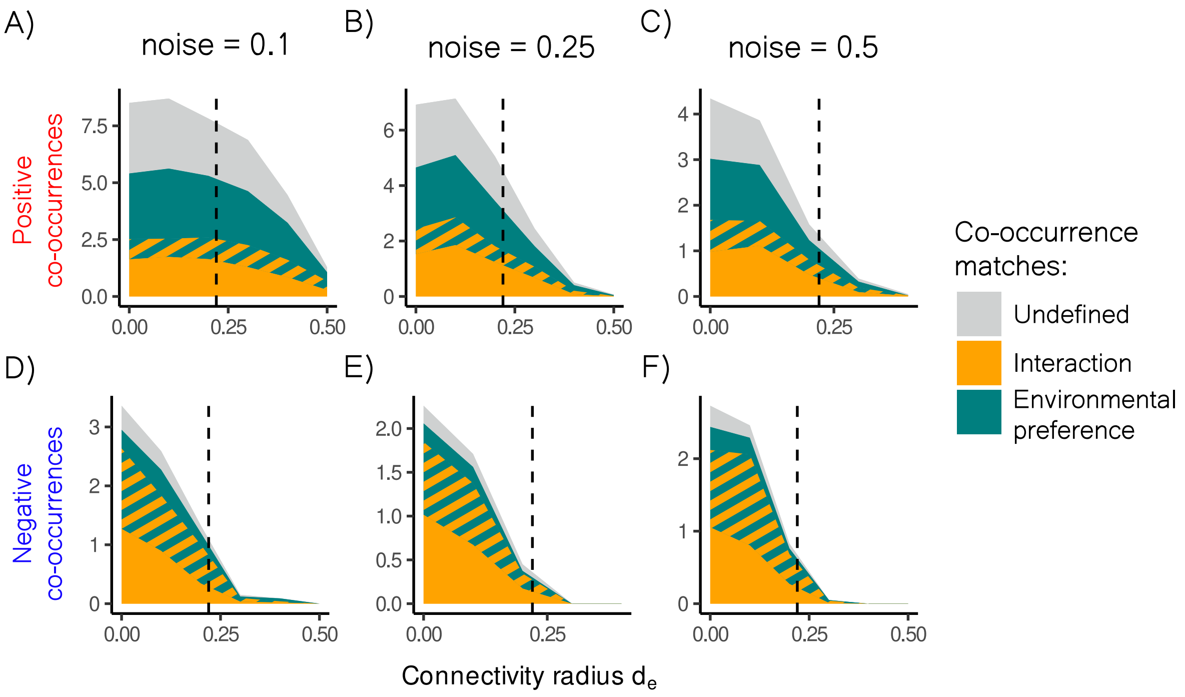

Figure S3: Three different levels of noise are added to simulations with the resource distribution of rs3. The amount of added noise is determined by the standard deviation of species abundances in habitats and a scaling factor $\xi$, which takes three values of 0.1, 0.25 and 0.5. Both observed positive and negative co-occurrences are shown with areas highlighted by they drivers they match. All values are averages of 100 runs.

##### Resource distribution

#### S4.1 Sampling scales

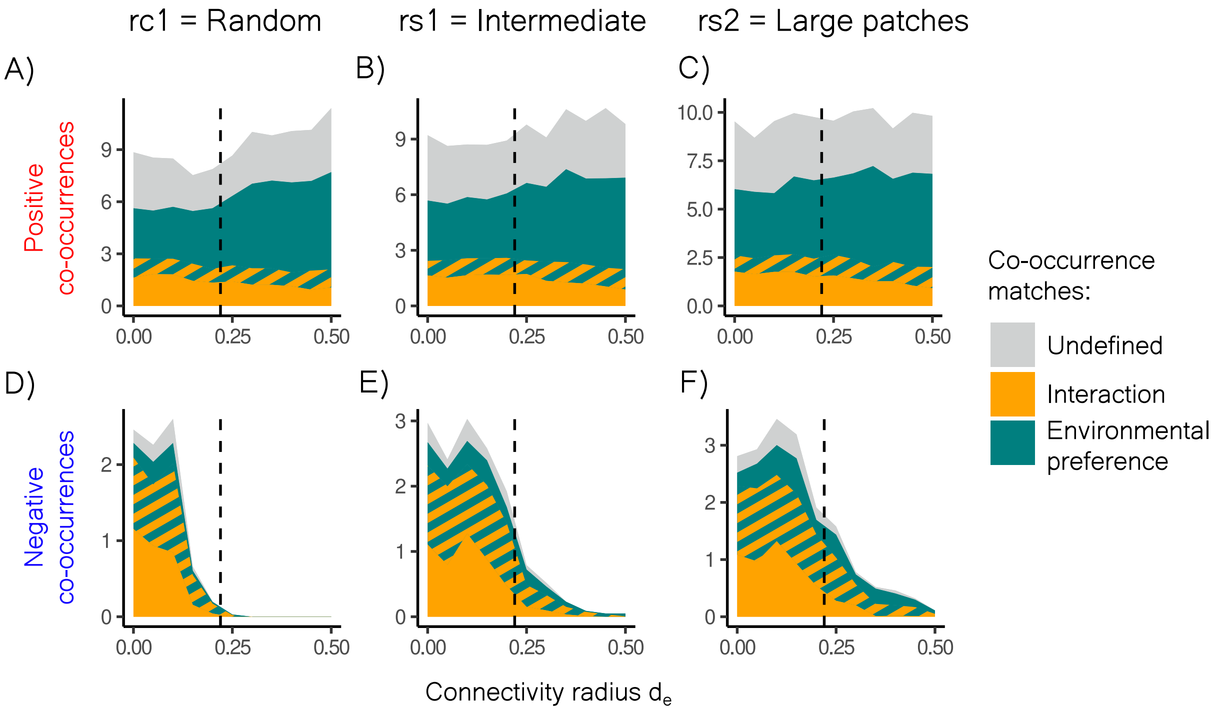

Figure S4: The effect of dispersal at different levels of $d_e$ is shown for the three resource distribution scenario not shown in the main manuscript. rc1 is the scenario in which resources show completely random distribution; rs1 shows intermediate distribution, while for rs2 resources are distributed in large patches. Both observed positive and negative co-occurrences are shown with areas highlighted by they drivers they match. All values are averages of 100 runs.

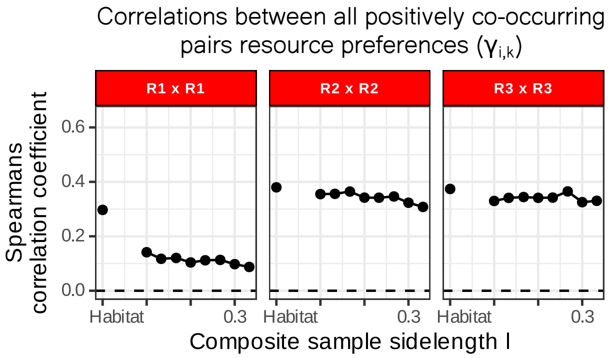

Figure S5: Spearman's correlation coefficients for positively co-occurring pairs resource preferences. All species pairs resource preference values $\gamma_{i,k}$ are correlated across all co-occurring pairs. Preferences for $R1$, $R2$ and $R3$ are shown. Correlations are calculated for all pairs found at different sampling volumes given different sample side lengths l. High values mean that there is a strong concordance with preference for a given resource and co-occurrence, while low values mean that preference for a given resource is less determinant of producing co-occurrence.

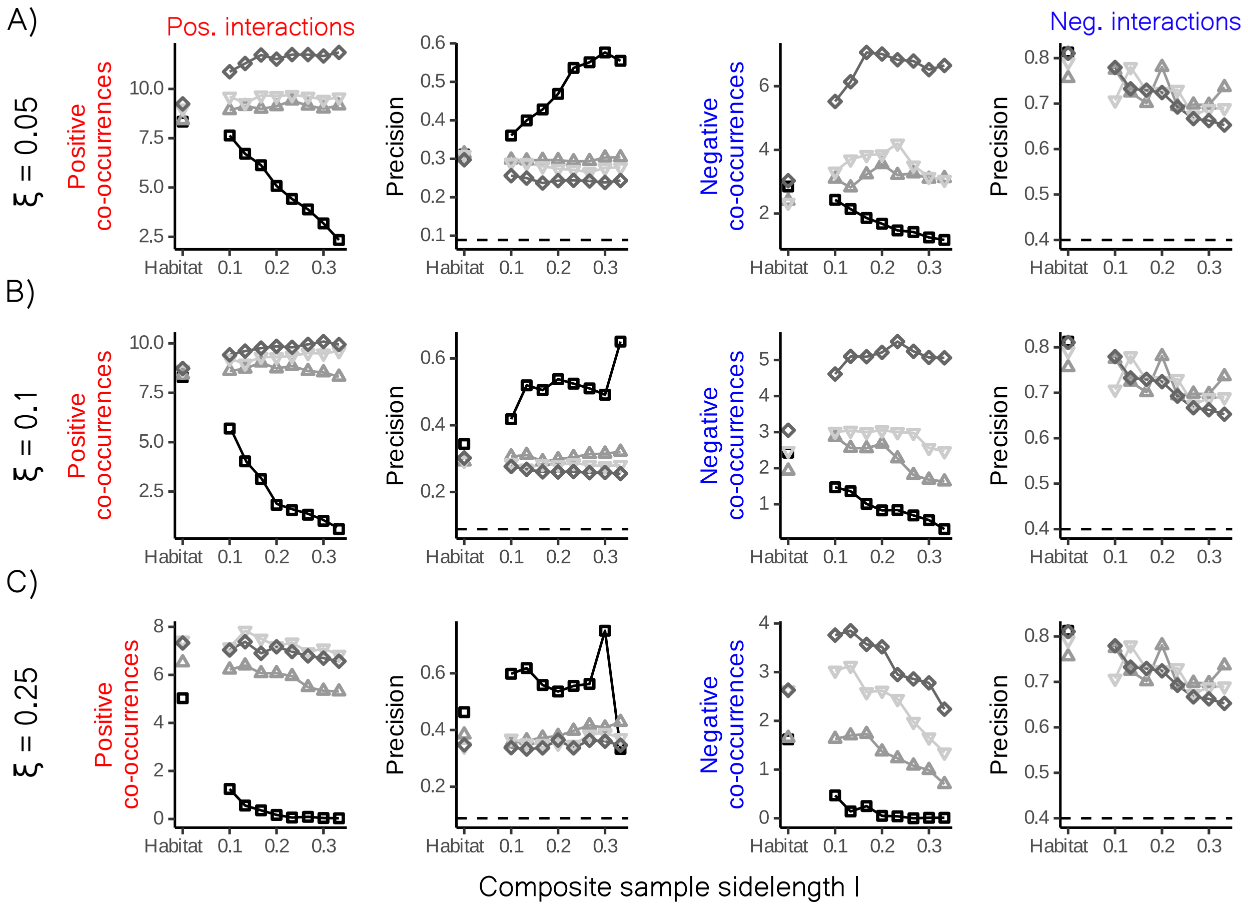

Figure S6: Three different levels of noise are added to simulations of all resource distribution scenarios. The amount of added noise is determined by the standard deviation of species abundances in habitats and a scaling factor $\xi$. Due to the large effect of low noise we the three values we depict are 0.05, 0.1 and 0.25 (rows). Columns show from left to right: the number of positive co-occurrences, the precision for detecting positive interactions, the number of negative co-occurrences and the precision for detecting negative interactions.

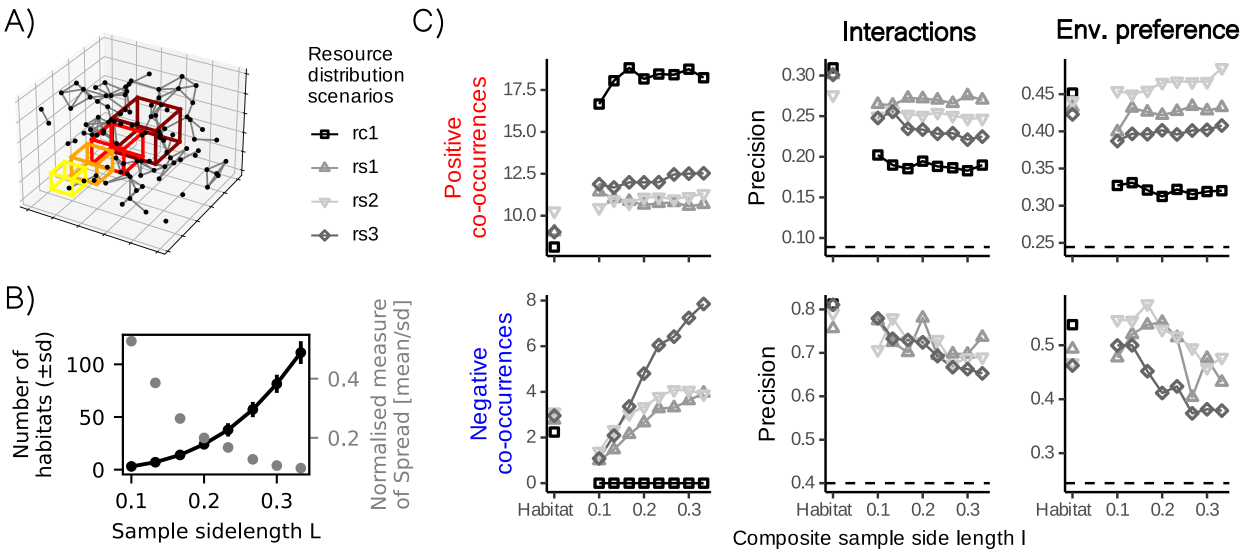

Figure S7: Composite sampling without fixing the number of habitats per samples and the influence of Simpsons paradox on the number of observed co-occurrences. A) The number of habitats per sample as well as the normalised measure of spread (standard deviation/mean) is shown at multiple scales of sampling. The number of B) positive and C) negative co-occurrences observed, when not correcting for the number of habitats sampled. Data is shown for all resource distribution scenarios and sampling scales. All data points are averages of 100 runs.
